# Molecular signature and target-specificity of inhibitory circuits formed by Martinotti cells in the mouse barrel cortex

**DOI:** 10.1101/2021.08.16.455953

**Authors:** Cristina Donato, Carolina Cabezas, Andrea Aguirre, Joana Lourenço, Marie-Claude Potier, Javier Zorrilla de San Martin, Alberto Bacci

## Abstract

In the neocortex, fast synaptic inhibition orchestrates both spontaneous and sensory-evoked activity. GABAergic interneurons (INs) inhibit pyramidal neurons (PNs) directly, modulating their output activity and thus contributing to balance cortical networks. Moreover, several IN subtypes also inhibit other INs, forming specific disinhibitory circuits, which play crucial roles in several cognitive functions. Here, we studied a homogeneous subpopulation of somatostatin (SST)-positive INs, the Martinotti cells (MCs) in layer 2/3 of the mouse barrel cortex (both sexes). MCs are a prominent IN subclass inhibiting the distal portion of PN apical dendrites, thus controlling dendrite electrogenesis and synaptic integration. Yet, it is poorly understood whether MCs inhibit other elements of the cortical circuits, and the connectivity properties with non-PN targets are unknown. We found that MCs have a strong preference for PN dendrites, but they also considerably connect with parvalbumin (PV)-positive, vasoactive intestinal peptide (VIP)-expressing and layer 1 (L1) INs. Remarkably, GABAergic synapses from MCs exhibited clear cell-type-specific short-term plasticity. Moreover, whereas the biophysical properties of MC-PN synapses were consistent with distal dendritic inhibition, MC-IN synapses exhibited characteristics of fast perisomatic inhibition. Finally, MC-PN connections used α5-containing GABA_A_Rs, but this subunit was not expressed by the other INs targeted by MCs. We reveal a specialized connectivity blueprint of MCs within different elements of superficial cortical layers. In addition, our results identify α5-GABA_A_Rs as the molecular fingerprint of MC-PN dendritic inhibition. This is of critical importance, given the role of α5-GABA_A_Rs in cognitive performance and their involvement in several brain diseases.

**Significance statement:** Martinotti cells (MCs) are a prominent subclass of SST-expressing GABAergic INs, specialized in controlling distal dendrites of PNs and taking part in several cognitive functions. Here we characterize the connectivity pattern of MCs with other INs in the superficial layers (L1 and L2/3) of the mouse barrel cortex. We found that the connectivity pattern of MCs with PNs as well as PV, VIP and L1 INs exhibit target-specific plasticity and biophysical properties. The stark specificity of α5-GABA_A_Rs at MC-PN synapses, and the lack or functional expression of this subunit by other cell types, define the molecular identity of MC-PN connections and the exclusive involvement of this outstanding inhibitory circuits in α5-dependent cognitive tasks.

## Introduction

In the neocortex, fast synaptic inhibition underlie important cognitive-relevant activity (Buzsáki, 2010; Isaacson and Scanziani, 2011). Neocortical inhibition is provided by GABAergic interneurons, which are highly heterogeneous and connect with both principal pyramidal neurons (PNs) and other inhibitory cells in a very stereotyped manner. Some interneurons, such as parvalbumin (PV)-expressing basket cells, innervate the perisomatic region of cortical PNs, and they thus provide a tight temporal control of PN spiking output and drive cognition-relevant fast network oscillations, especially in the β-γ-frequency range (20-100 Hz)(Bartos et al., 2007; Buzsáki and Wang, 2012).

Conversely, other interneuron types, such as those expressing the neuropeptide somatostatin (SST), were shown to target dendrites of PNs, thereby controlling dendritic electrogenesis, non-linear integration and glutamatergic synaptic input (Wang et al., 2004; Lovett-Barron et al., 2012; Wilson et al., 2012; Schulz et al., 2018). In sensory cortices, SST interneurons were shown to be involved in lateral inhibition, playing a major role in key sensory computations, such as surround suppression (Kapfer et al., 2007; Silberberg and Markram, 2007; Berger et al., 2009; Adesnik and Scanziani, 2010; Adesnik et al., 2012). Moreover, SST-operated dendritic inhibition was shown to encode fear memory and affective behavior in prefrontal cortex (Xu et al., 2013; Scheggia et al., 2019; Clem and Cummings, 2020).

SST INs were proposed to be the source of a profuse ‘blanket’ of inhibition due to their dense connectivity with PNs (Fino and Yuste, 2011). However, this view neglects the diversity of SST-positive INs (Gouwens et al., 2020), and the fact that they preferentially contact specific PN subclasses (Hilscher et al., 2016) as well as other inhibitory neurons (Pfeffer et al., 2013; Tremblay et al., 2016). In particular, SST interneurons can be classified as Martinotti cells (MCs) and non-Martinotti cells, which exhibit differential connectivity patterns as well as specific molecular profiles (Wang et al., 2004; Ma et al., 2006; Tremblay et al., 2016; Yavorska and Wehr, 2016; Paul et al., 2017; Scala et al., 2019). In particular, MCs exhibit a well-defined axonal morphology, as they project their axons to layer 1, where they extensively inhibit the most distal dendritic tufts of PNs (Wang et al., 2004; Ma et al., 2006; Kapfer et al., 2007; Silberberg and Markram, 2007; Tremblay et al., 2016). Functionally, MCs are efficiently recruited by local PNs with loose-coupled, strongly facilitating synapses (Reyes et al., 1998; Wang et al., 2004; Kapfer et al., 2007; Silberberg and Markram, 2007), and are quasi-preferentially inhibited by vasoactive intestinal peptide (VIP)-expressing GABAergic interneurons (Pfeffer et al., 2013; Karnani et al., 2016; Tremblay et al., 2016; Walker et al., 2016). Finally, MCs form synapses with other elements of the cortical circuit, namely other inhibitory interneurons (Ma et al., 2006; Pfeffer et al., 2013). However, the actual extent and biophysical properties of these disinhibitory circuits are unknown and/or generalized over SST-expressing MCs and nMCs (Pfeffer et al., 2013).

Importantly, MC-PN inhibitory synapses were shown to use the α5-containing GABAAR (α5-GABA_A_Rs) (Ali and Thomson, 2008; Zorrilla de San Martin et al., 2020). Similarly, the hippocampal counterparts of MCs, the oriens-lacunoso moleculare (O-LM) interneurons express functional α5-GABA_A_Rs (Schulz et al., 2018). This prompts the question whether GABAergic synapses formed by MCs onto other elements of the cortical circuit use this specific subunit of GABA_A_Rs. Understanding the actual synaptic circuits relying on the α5 subunit has important clinical implications. Indeed, α5-GABA_A_Rs were indicated as a prominent target for therapeutic interventions for cognitive dysfunctions in Down syndrome (Braudeau et al., 2011; Duchon et al., 2019; Schulz et al., 2019; Zorrilla de San Martin et al., 2020), depression (Zanos et al., 2017), anesthesia-induced memory impairment (Zurek et al., 2014) and schizophrenia (Duncan et al., 2010; Gill and Grace, 2014).

Here we investigated the connectivity blueprint of MCs in the superficial layers of the mouse barrel cortex. We found that, in addition to the known connectivity with PN distal dendrites, MCs connect extensively also with PV, VIP and L1 INs, but not with other MCs. Interestingly, GABAergic synapses formed by MCs exhibited clear target specificity of short-term plasticity. Finally, dendritic inhibition using α5-GABA_A_Rs is a peculiarity of MC-PN synapses, as unitary responses from MCs to other INs exhibited fast (<1ms) rise-time, and they were not modulated by a α5 negative allosteric modulator (NAM).

Altogether, these results indicate the molecular, connectivity and biophysical fingerprint used by MCs for inhibitory synapses that they make with PNs and other elements of the cortical circuit.

## Materials and Methods

### Animals

Experimental procedures followed national and European (2010/63/EU) guidelines and have been approved by the author’s institutional review boards and national authorities (APAFIS #2599). All efforts were made to minimize suffering and reduce the number of animals. Mice used in this study were of both sexes. In order to identify GABAergic transmission from different INs we used several mouse models. To record from PV INs we initially used *Pvalb*-cre mice (Jackson Laboratory, Stock Number: 008069), crossed with a mouse line, which expresses a *loxP*-flanked STOP cassette and giving robust tdTomato fluorescence following Cre-mediated recombination (Jackson Laboratory Stock Number 007909). In the experiments illustrated in Figs. 2,3,5 and 6, we used PValbTomato mouse line (Kaiser et al., 2016, Jackson Stock# 27395), a line that expresses TdTomato fluorescent protein specifically in PV INs. To record from MCs, we used GAD-67 GFP X98 mice (Ma et al., 2006), herein defined as X98. These mice express EGFP in a specific subset of GABAergic cells (Jackson Laboratory Stock# 006340). To perform simultaneous recordings from MCs and PV INs we crossed X98 mice with PvAlb-tdtomato. Furthermore, in order to record from synaptically connected VIP INs and MCs we crossed VIP-Cre mice (Jackson Laboratory Stock #010908) with X98 mice and infected newborns with viral vectors carrying the genes of either ChR2 or TdTomato (see below details of different viral infections).

### In Vitro Slice Preparation and Electrophysiology

Coronal slices (300-350 μm thick) from somatosensory cortex were obtained from 18- to 25-d-old mice. Animals were deeply anesthetized with isoflurane and decapitated. Brains were quickly removed and immersed in “cutting” solution (4°C) containing the following (in mM): 126 choline, 11 glucose, 26 NaHCO_3_, 2.5 KCl, 1.25 NaH_2_PO_4_, 7 MgSO_4_ and 0.5 CaCl_2_ (equilibrated with 95-5% O_2_-CO_2_, respectively). Slices were cut with a vibratome (Leica) in the same cutting solution and then incubated in oxygenated artificial cerebrospinal fluid (aCSF) containing the following (in mM): 126 NaCl, 2.5 KCl, 2 CaCl_2_, 1 MgSO_4_, 1.25 mM NaH_2_PO_4_, 26 mM NaHCO_3_, and 16 mM glucose (pH 7.4), initially at 34°C for 30 min, and subsequently at room temperature until transfer to the recording chamber. Recordings were obtained at 32-34°C. Whole-cell voltage-clamp recordings were performed in from layer (L)2/3 PNs, MCs, PV, VIP INs and L1 INs of the primary somatosensory cortex. PNs were visually identified using infrared video microscopy by their large somata and pia-oriented apical dendrites. L1 INs were also visually identified with transmitted light only as they are the only cell type with the soma present in L1. MCs (labeled with GFP, see Fig 1), VIP INs and PV INs (labeled with TdTomato), were identified using LED illumination (blue, λ=470nm, green λ=530nm, OptoLED system, Cairn Research, Faversham, UK) coupled to epifluorescent optical pathway of the microscope. Single or double voltage-clamp whole-cell recordings were made with borosilicate glass capillaries (with a tip resistance of 2–4 MΩ) filled with different intracellular solutions depending of the experiment. For unitary inhibitory postsynaptic currents (uIPSCs) the intracellular solution contained (in mM): 70 K-gluconate, 70 KCl, 10 HEPES, 1 EGTA, 2 MgCl_2_, 4 Mg-ATP, 0.3 Na-GTP, pH adjusted to 7.2 with KOH, 280–300 mOsm or 145 CsCl, 4.6 MgCl_2_, 10 HEPES, 1 EGTA, 0.1 CaCl_2_, 4 Na-ATP, 0.4 Na-GTP, pH adjusted to 7.2 with CsOH, 280–300 mOsm. To confirm the GABAergic nature of uIPSCs, gabazine (10 μM) was added to the aCSF at the end in some experiments. For tonic inhibition experiments, GABA (5 μM) was added to the aCSF. To record unitary excitatory postsynaptic currents (uEPSCs) from INs, a low chloride intracellular solution was used and DNQX was omitted in the aCSF superfusate. In these experiments, the intracellular solution had the following composition (in mM): 150 K-gluconate, 4.6 MgCl_2_, 10 HEPES, 1 EGTA, 0.1 CaCl_2_, 4 Na-ATP, 0.4 Na-GTP, pH adjusted to 7.2 with KOH, 280–300 mOsm. In voltage-clamp experiments, access resistance was on average <15 MΩ and monitored throughout the experiment. Recordings were discarded from analysis if the resistance changed by >20% over the course of the experiment. Unitary synaptic responses were elicited in voltage-clamp mode by brief somatic depolarizing. A train of 5 presynaptic spikes at 50 Hz was applied to infer short-term plasticity of synaptic responses. The paired pulse ratio (PPR) was obtained as the peak amplitude of the second uEPSC divided by that of the first. In order to isolate GABAA-receptor-mediated currents, DNQX (10 μM) was present in the superfusate of all experiments, unless otherwise indicated.

**Figure 1:**
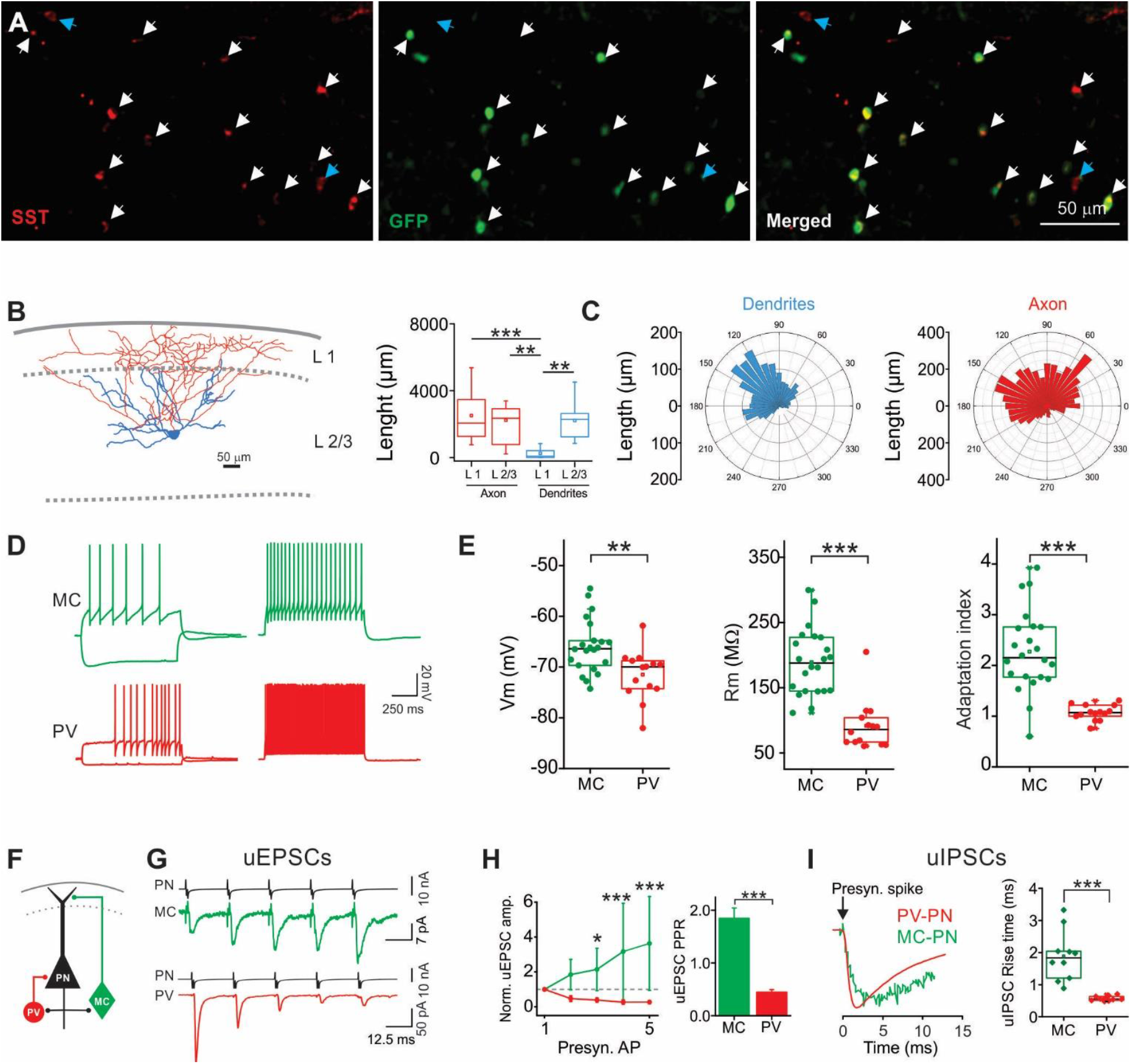
GFP-positive neurons in X98 mice are SST-expressing MCs. **A:** Top: Epifluorescence micrograph of a dual immune-staining against SST (red) and GFP (green) in X98 coronal somatosensory slices. White arrows: GFP and SST co-localization; blue arrows: cells expressing SST only. **B:** Left: representative morphological reconstruction of a GFP-positive neuron filled with biocytin. Blue: dendrites; red: axons. Right: Population data of axon (red) and dendrite (blue) lengths distribution in L1 and L2/3 (n=11). **C:** Axonal (red) and dendritic polar plots of the cell of B. **D:** Representative current-clamp recordings from a GFP-expressing interneuron in X98 mice (green) and a PV cell (red). X98 GFP cells display a characteristic sag in response to hyperpolarizing current injection and a highly adapting firing behavior. Conversely, PV-cells show fast-spiking patterns in response to depolarizing current injections. **E:** Summary graphs of resting membrane potential (left), membrane resistance (middle) and adaptation index (right) in PV interneurons (n=14) and MCs (n=22). **F:** Schematic of mutually connected MC-PN and PV-PN pairs. **G:** Representative averaged voltage clamp trace of unitary EPSCs stimulated by 5 action potentials at 40Hz in a PN and recorded in a GFP-positive cell from a X98 mouse (green, upper panel), and in a PV-cell (bottom panel). **H:** Left panel: pooled normalized amplitudes of uEPSC evoked with a 50Hz, 5 AP train. Right: population plot of paired-pulse ratio (PPR) of X98 GFP (n=20, green) and PV-INs (n=11, red). **I:** Left: overlapped representative uIPSCs elicited by MCs (green) and PV-INs (red) recorded PNs. Right: population plot of the uIPSC mean rise time from MC to PN (green) and PV to PN (red) synapses. * p<0.05, ** p<0.01, *** p<0.001.

Signals were amplified, using a Multiclamp 700B patch-clamp amplifier (Molecular Devices, San Jose, CA), sampled at 20-100 kHz and low-pass filtered at 4 KHz (for voltage clamp experiments) and 10 KHz (for current clamp experiments). All drugs were obtained from Tocris Cookson (Bristol, UK) or Sigma (Bristol, UK). α5IA, (3-(5-methylisoxazol-3-yl)-6-[(1-methyl-1,2,3-triazol-4-yl)methyloxy]-1, 2, 4-triazolo[3, 4-a]phthalazine) also named L-822179 was synthesized by Orga-Link SARL (Magny-les-Hameaux, France) according to Sternfeld et al. (2004) as in Braudeau et al. (2011). The hydrochloride salt was solubilized in DMSO at a concentration of 1mM and then diluted in the appropriate buffer.

### Virus-Mediated Gene Delivery and Optogenetics

To study MC-VIP and VIP-MC synapses we first crossed crossed VIPcre with X98 mice and injected 300 nL of a solution containing adeno-associated viral (AAV) particles into the somatosensory cortex of ice–anesthetized pups (P0–3) to selectively express TdTomato or Channelrhodopsin-2 (ChR2) in VIP INs. Injections were made with a beveled glass pipette 300 μm deep in the somatosensory cortex through intact skin and skull. We then delivered the solution containing the AAVs using a Nanoliter 2000 Injector (WPI Inc., USA). The pipette was left in place for an additional 30 s, before it was retracted. The AAVs expressed floxed ChR2 or TdTomato (AAV9.EF1.dflox.hChR2(H134R)-mCherry.WPRE.hGH; Addgene #20297 and pAAV-FLEX-tdTomato; Addgene #28306, respectively) purchased from the Penn Vector Core (University of Pennsylvania). At the end of the procedure, pups were returned to their mother. ChR2 activation was obtained by brief (0.5-2 ms) LED light pulses on cortical slices (λ = 470 nm). Experiments were performed using a 60X water immersion lens. Light-evoked responses were recorded in L 2/3 MCs and were completely abolished by gabazine (not shown).

### Data analysis

Experiments on firing dynamics, tonic currents and unitary paired recordings were analyzed with Clampfit (Molecular Devices), Origin (Microcal) and custom-made scripts in Matlab (the Mathworks). Spontaneous synaptic events were detected using custom written software (Wdetecta, courtesy J. R. Huguenard, Stanford University https://hlab.stanford.edu/wdetecta.php) based on an algorithm that calculates the derivative of the current trace to find events that cross a certain defined threshold. Amplitude and rise times of the events were then binned and sorted, using other custom written routines (courtesy J. R. Huguenard, Stanford University).

The peak-to-baseline decay phase of uIPSCs was fitted by the following double exponential function:

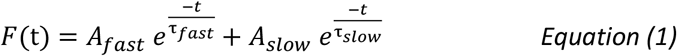

where *A_fast_* and *A_slow_* are the fast and slow amplitude components, and *τ_fast_* and *τ_slow_* are the fast and slow decay time constants, respectively. The weighted decay time constant (τ_d,w_) was calculated using the following equation:

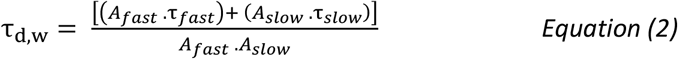

The adaptation index was calculated as the last/first inter-spike interval ratio following a train of spikes induced by injection of a depolarizing step of current. Passive properties as well as synaptic currents were analyzed with Clampfit and custom-made scripts in MATLAB (Mathworks). Both unitary and light-induced IPSCs were averaged across at least 20 sweeps for each condition examined. Results are presented as means ± SEM unless otherwise stated.

### Morphological reconstruction

To reconstruct and quantify anatomical features of different cortical neurons, biocytin (Sigma) was included in the intracellular solution at a high concentration (10mg/mL), which required extensive sonication. To avoid excessive degradation of fragile molecules such as ATP, sonication was performed in an ice bath. The intracellular solution was then filtered twice to prevent the presence of undissolved lumps of biocytin in the patch pipette. Recordings lasted for at least 30 min. During that time, access resistance was continuously monitored throughout the experiment. At the end of recordings, the patch pipette was removed carefully to obtain an outside-out patch in order to reseal the cell properly. The slice was then left in the recording chamber for a further 5-10 min to allow biocytin diffusion. Slices were then fixed with 4% paraformaldehyde in phosphate buffer saline (PBS, Sigma) for at least 48h. Following fixation, slices were incubated with the avidin-biotin complex (Vector Labs) and a high concentration of detergent (Triton-X100, 5%) for at least two days before staining with 3,3’-Diaminobenzidine (DAB, AbCam). Cells were then reconstructed, and cortical layers delimited using Neurolucida (MBF Bioscience). Neuronal reconstructions were aligned to a mouse atlas from the Allen Institute. By using Neurolucida Explorer, we analyzed the length of axons and dendrites of MCs in L2/3 and L1 of somatosensory cortex. Data were exported and analyzed in OriginPro 2016 (OriginLab Corporation).

### Immunofluorescence

Slices used for electrophysiology experiments were fixed overnight in 4% paraformaldehyde in phosphate buffered saline (PBS, pH 7.4) at 4°C. Slices were then rinsed three times at room temperature (10 min each time) in PBS and incubated overnight at 4°C in PB with 0.3% Triton X-1000, 0.1% normal goat serum (NGS), anti-GFP antibody (host: rabbit, 1∶400, AB3080, Millipore) and/or anti-SST antibody (host: mouse, 1:250, G10 sc-55565, Santa Cruz) and/or anti-DsRed (host: rabbit, 1/500, #632496 Takara Bio Clontech). Slices were then rinsed three times in PBS (10 min each) at room temperature and incubated with goat-hosted secondary antibodies coupled to different fluorophores: Alexa 488 (1:500, A11034, Life technologies) and Alexa 633 (1∶500, A21052, Life technologies) for 2 h at room temperature. Slices were then rinsed three times in PBS (10 min each) at room temperature and mounted with Fluoromount. Immunofluorescence was then images were acquired with a confocal microscope (Leica SP8).

Parvalbumin, SST and GFP staining on X98 mice were also performed on 50 μm-thick slices. Briefly, mice were perfused with 0.9% NaCl solution containing Heparin and 4% paraformaldehyde (PFA). Brains were cryo-protected by placing them overnight in 30% sucrose solution and then frozen in Isopentane at a temperature <−50°C. Brains were sliced with a freezing microtome (ThermoFisher HM450). Permeabilization in a blocking solution of PBT with 0.3% Triton and 10% Normal Goat Serum was done at room temperature for 2 hr. Slices were then incubated overnight (4°C) in the same blocking solution containing the primary rabbit anti-PV antibody (1:1000; Thermo Scientific, PA1-933) and mouse anti-GFP antibody (1:500; Milipore MAB3580). Slices were then rinsed three times in PBS (10 min each) at room temperature and incubated with goat anti-rabbit and a goat anti-mouse antibody (1:500; Jackson IR) coupled to Alexa-488 or 633 for 3.5 hr at room temperature. Slices were then rinsed three times in PBS (10 min each) at room temperature and coverslipped in mounting medium (Fluoromount, Sigma Aldrich F4680). Immunofluorescence images were acquired with a confocal (Leica SP8) or epifluorescence (Zeiss Apotome 3) microscope.

### Statistical Analysis

All statistical analysis were performed in Origin (Microcal). Normality of the data was systematically assessed (Shapiro-Wilk normality test). Normal distributions were statistically compared using Paired t-Test or Two-sample t-Test. When data distributions were not normal or n was small, non-parametric tests were performed (Mann-Whitney, Wilcoxon Signed Ranks Test). Two-way ANOVA tests were followed by Bonferroni’s multiple comparison post hoc. Differences were considered significant if p <0.05 (*p<0.05, **p<0.01, ***p<0.001).

## Results

### The X98 mouse is a reliable model to specifically study L2/3 Martinotti cells

Despite being broadly classified as dendrite-targeting INs, SST-expressing cells exhibit significant electrophysiological, anatomical, connectivity and molecular heterogeneity (Ma et al., 2006; Paul et al., 2017; Naka et al., 2019). In order to specifically study the connectivity of L2/3 MCs we searched for a suitable mouse line. X98 mice express GFP predominantly in cortical layers (L) 5B and 6, and, to a lesser extent, in L2/3 (Ma et al., 2006). A detailed characterization of these mice showed that GFP is specifically expressed in L5 MCs (Ma et al., 2006). However, although there is prominent fluorescence in L2/3, GFP-expressing cells in this cortical layer were not analyzed. Therefore, we first set out to confirm that GFP-expressing cells in L2/3 belong to the specific SST-positive interneuron subtype defined as the MCs. We performed immunofluorescence staining on microtome-cut sections from X98 coronal somatosensory slices of 18-25-days-old mice and showed that all GFP-expressing cells also expressed SST while some SST-positive cells did not express GFP (Fig. 1 A). In another series of experiments, several GFP-expressing neurons were filled with biocytin during whole-cell recordings and their morphology was traced to assess somato-dendritic and axonal morphology. Axons of L2/3 GFP-expressing neurons were systematically oriented towards superficial layers and consistently reached L1 where they were profusely branched (red tracing in Fig. 1 B and C; p=4.4e^-4^, One-way ANOVA followed by Bonferroni post-hoc test, F=7.4716,, n=11 reconstructed GFP-positive neurons). Conversely, GFP-expressing neurons dendrites were mostly located in L2/3 without reaching L1 (blue tracing in Fig. 1 B and C). We then assessed the excitability and passive properties of GFP-expressing neurons (n=22) and compared their firing pattern with that of PV INs, the most abundant and perhaps best characterized GABAergic neuronal subtype (Fig 1 D). As previously described, the majority of the GFP-positive cells in X98 mice displayed a characteristic sag in response to hyperpolarizing current injection and a highly adapting firing behavior when depolarizing currents triggered repetitive spiking (Fig 1 D). Conversely, PV INs displayed fast-spiking, non-adapting pattern in response to depolarizing currents (adaptation index: 2.27 ± 0.17 and 1.07 ± 0.04 for GFP-expressing and PV INs, respectively; p=1.1e^-5^, Mann-Whitney U test; Fig. 1 D), more hyperpolarized resting membrane potential (Vm: −66 ± 1 and −71 ± 1 mV for GFP-expressing and PV INs, respectively; p=0.0017, unpaired T test) and lower input resistance (Ri: 189 ± 11 and 92 ± 10 MΩ for GFP-expressing and PV INs, respectively; p=8.1e^-6^, Mann-Whitney U test; Fig. 1 E).

We then assessed the biophysical and pharmacological traits of synaptic transmission that distinguish MCs from other INs. We analyzed unitary glutamatergic, excitatory and GABAergic, inhibitory currents (uEPSCs and uIPSCs, correspondingly) onto and from putative MCs in MC-PN connected pairs. One hallmark of MC connectivity is the strongly facilitating glutamatergic synaptic responses evoked upon PN action potentials (Wang et al., 2004; Kapfer et al., 2007; Silberberg and Markram, 2007). Accordingly, we found that unitary excitatory inputs from PNs to putative MCs in X98 mice were invariably facilitating while uEPSCs onto PV INs were depressing (paired pulse ratio: 1.8 ± 0.2 for GFP-expressing neurons and 0.4 ± 0.1 for PV INs; p=1.8e^-6^, Mann-Whitney U test; Fig 1 F-H). Finally, we analyzed the kinetics of uIPSCs elicited by MCs and PV-cells in L2/3 PNs known to have distinctive kinetics (Silberberg et al., 2007). We found that uIPSCs evoked from MCs had significantly slower rise times as compared to PV INs (Rise time: 1.89 ± 0.25 ms for GFP-expressing neurons and 0.57 ± 0.02 for PV INs; p=2.2e^-5^, unpaired T test; Fig. 1 I), consistent with characteristic MC-mediated dendritic uIPSCs.

Altogether, these results indicate that GFP-expressing neurons in L2/3 of the somatosensory cortex of X98 mice are a homogeneous subgroup of MCs as they exhibit typical anatomical, intrinsic excitability and synaptic features of MCs. Furthermore, this subgroup can be readily distinguished from the most abundant GABAergic PV INs.

### Martinotti Cells display target-specific synaptic properties

In addition to PNs, SST INs were shown to contact other inhibitory neurons of the cortical microcircuits including VIP, PV and L1 INs (Pfeffer et al., 2013; Tremblay et al., 2016). However, it remains unknown whether this is true for MCs and whether these connections exhibit the biophysical and pharmacological properties observed in the inhibitory MC-PN synapse. In order to address this question, we systematically evoked uIPSCs from specific synapses formed between MCs and other INs using dual patch recordings in brain slices. To measure and compare evoked uIPSCs from pairs between MCs and other INs, we used brain slices containing differently labeled IN subtypes. For MC-PV synapses, we crossed X98 mice with *Pvalb-*tdTomato mice. For MC-L1 synapses, we used X98 mice and L1 INs were identified by their localization in L1. Finally, to record uIPSCs from MC-VIP cell pairs we crossed VIPcre with X98 mice. Mouse pups (P1-3) were then subjected to intracerebral injections of flexed AAV particles coding for tdTomato. We could thus obtain mice, in which MCs and VIP cells were simultaneously labeled with GFP and tdTomato, respectively.

We found significant connectivity rates between MCs and PV INs (13 connected out of 85 recorded pairs), between MCs and L1 INs (11 connected out of 80 recorded pairs) and between MC and VIP INs (9 connected out of 45 recorded pairs; Fig. 2 B). Yet, the connectivity rate between MCs and these IN types was much lower than functional connections with PNs (30 connected out of 57 recorded pairs). Conversely, we did not find functional synaptic transmission between MCs (0 out of 10, connected/recorded pairs; Fig 2 B). Amplitudes of evoked uIPSCs were also largely variable between and within synapses. uIPSC amplitudes were consistently larger in MC-IN than in MC-PN synapses (uIPSC amplitudes: 10 ± 1; 36 ± 4; 72 ± 32; 172 ± 70 pA; MC-PN, -PV, L1 and VIP INs, respectively; Kruskal Wallis followed by Mann-Withney with Bonferroni’s correction; n = 10, 7, 5, 6, respectively).

**Figure 2:**
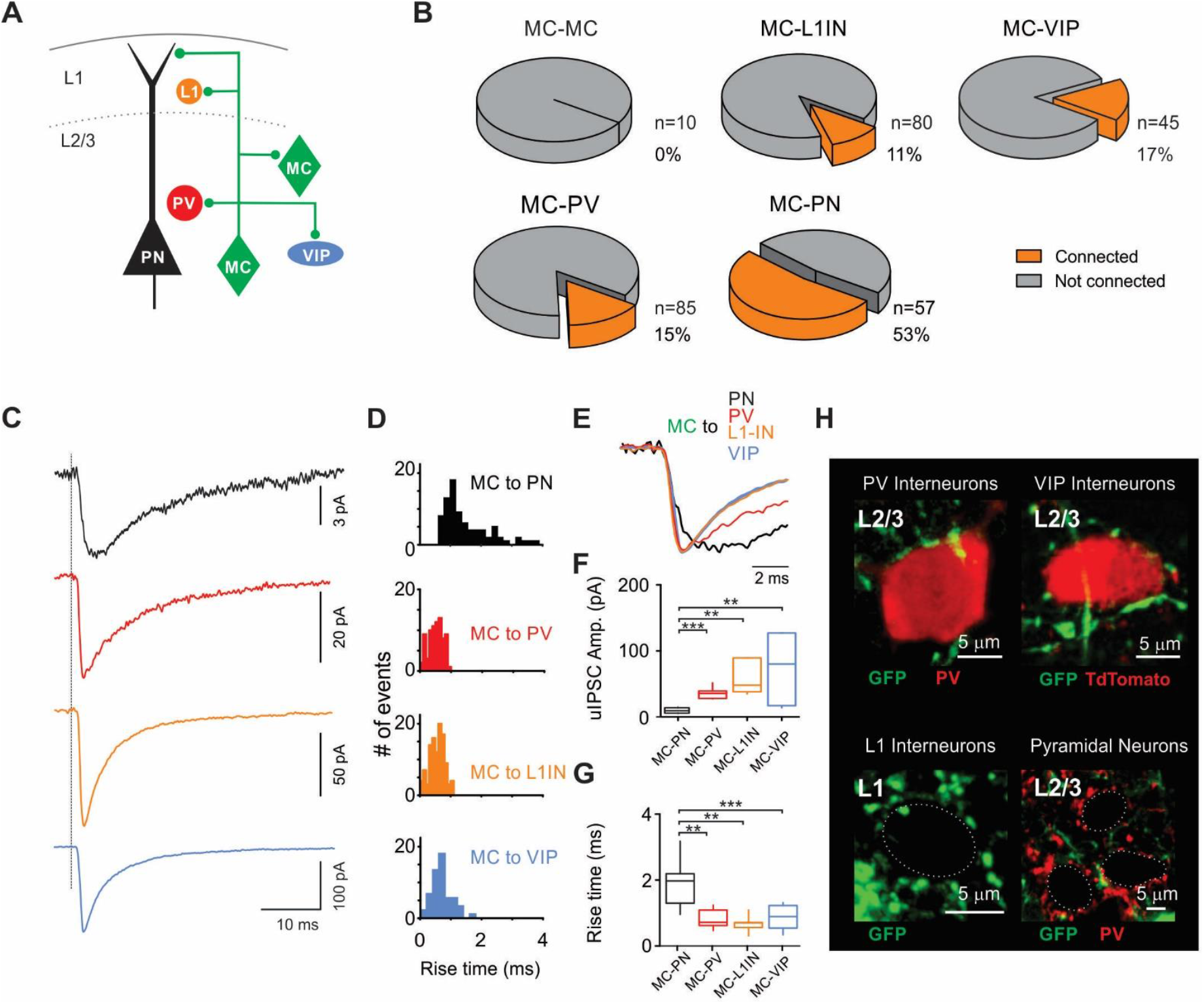
Diversity of MC synaptic contacts onto different neuronal types in the L2/3 of somatosensory cortex. **A:** Schematic representation of the tested inhibitory connections involving MCs. **B:** Pie charts illustrating the connectivity rates. **C:** Representative voltage-clamp uIPSCs average traces from MC-PN (black), MC-PV-IN (red) and MC-L1 IN (orange), VIP-INs (blue). Gray dotted line represents the time of the peak of presynaptic action potentials. **D:** Representative distributions of uIPSCs Rt recorded from individual MC-PN (top, black), MC-PV-IN (middle, red) and MC-L1 (bottom, orange) connections. **E:** uIPSC (same as in C) normalized to the peak. **F:** Mean uIPSC amplitudes. **G:** Population plot of the mean uIPSCs Rts from individual MC-PN, MC-PV, MC-L1, PV-PN and MC-VIP recorded connections. **H:** Confocal micrographs illustrating immunolabelling of: GFP-expressing MC puncta and PV (top left), GFP-expressing MC puncta and TdTomato-expressing VIP-IN (top right), GFP-expressing MC puncta around L1 IN soma (bottom left), GFP-expressing MC puncta and PV-positive synaptic baskets formed around L2/3 PN somas (bottom right). ** p<0.01, *** p<0.001.

GABAergic synapses formed by MCs to PNs are slow due to their distal dendritic location and consequent electrotonic filtering. To further explore whether synaptic contacts made by MCs onto other circuit elements followed a similar pattern, we compared the kinetics of uIPSCs elicited by MCs onto PNs, PV-, L1-and VIP-INs (Fig. 2 C-F). Rise time (Rt) of MC-PN uIPSCs were significantly slower than those recorded from MC-PV, MC-L1 and MC-VIP-IN pairs (1.89 ± 0.25; 0.73 ± 0.10; 0.63 ± 0.13; 0.80 ± 0.15 ms, respectively; p=1.5e^-4^, one-way ANOVA; n = 10, 7, 5, 6, respectively; Fig 2 C-F). Rise times of uIPSCs recorded from connected pairs between MCs and PV, VIP and L1 INs were not significantly different (one-way ANOVA followed by Bonferroni post hoc test).

One possible explanation for the different uIPSC kinetics observed at inhibitory synapses made by MCs could be that synaptic inputs located in the somatic/perisomatic region are less filtered than those located in distal dendrites. Therefore, we took advantage of the fluorescent labeling strategies to analyze putative contacts between GFP-positive MC axons and the somatic compartment of PV-, L1- or VIP-INs. Analysis of confocal images revealed GFP-positive MC axons and bona fide boutons juxtaposed to PV-IN somas and to VIP-INs in L2/3 (Fig 2H). Moreover, in L1 we also found L1 IN somas circled by GFP-positive MC bona fide puncta (Fig 2 G). Conversely, PN somas in L2/3 were profusely surrounded by PV-positive puncta but almost no GFP-positive puncta from MCs. Although indirect, this evidence suggests that MCs make axo-somatic connections onto other INs while they make exclusively axo-dendritic connections onto PNs.

We then analyzed short-term synaptic plasticity (STP) at all unitary connections made by MCs with different postsynaptic targets (Fig. 3A), in response to trains of 5 action potentials at 50 Hz. We found that short-term plasticity profiles depended on the postsynaptic target. Indeed, GABAergic transmission at MC-PN and MC-L1 IN synapses were strongly depressing. In contrast, MC-PV uIPSCs did not vary during the stimulus train, and MC-VIP synapse exhibited a significant facilitating profile (Fig. 3B-C). When compared with MC-PN connections, STP at MC-L1 IN synapses were not significantly different. However, STP of MC-PV and MC–VIP IN synapses was significantly different than MC-PN connections (repeated measures, two-way ANOVA followed by Bonferroni post hoc test; F=24.1516, p=7.34e^-5^, n=5, 7, 5 and 7 synapses for MC-VIP, -PV, -L1 and –PN, respectively; Fig. 3 B-C).

**Figure 3:**
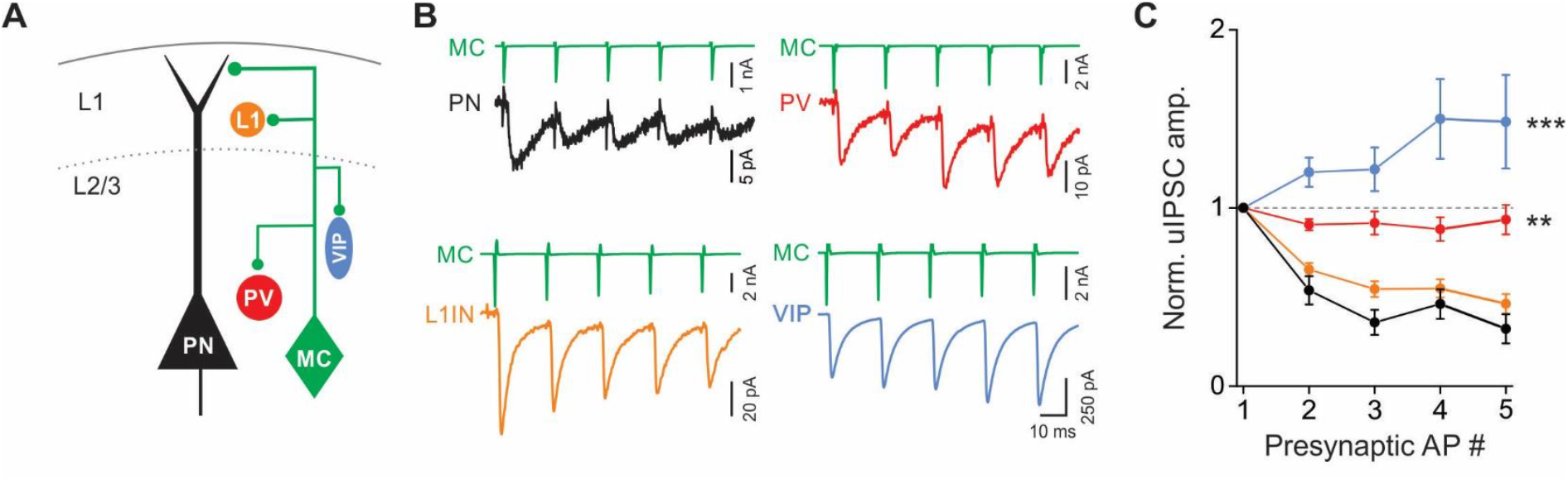
Plasticity of MC-mediated synaptic inhibition in L2/3 of somatosensory cortex. **A:** Schematic representation of the tested inhibitory circuits involving MCs. **B:** Representative voltage-clamp averaged traces of uIPSCs from MCs onto PNs (black), PV-cells (red), L1-Ins (orange) and VIP-Ins (blue). **C:** Normalized uIPSC amplitude elicited by MCs onto different elements of the L2/3 inhibitory circuit. Inhibition of MCs onto PN (black) and L1-INs (orange) is strongly depressing whereas connections onto VIP (blue) are facilitating. A slight facilitation occurs at MC-PV synapses.

Altogether, these results indicate that L2/3 MCs preferentially contact PNs, to a lesser, albeit non-negligible, extent PV, VIP and L1 INs, and avoid connecting between themselves. MC-dendrite targeting is specific for connections with PNs. Surprisingly, short-term plasticity at MC-synapses exhibit clear target specificity.

### α5-GABA_A_Rs define MC-PN synapses in L2/3 of mouse somatosensory cortex

MC-PN synapse has been shown to be mediated by GABA_A_Rs containing the α5 subunit in the rat somatosensory cortex (Ali and Thomson, 2008), in the mouse prefrontal cortex (Zorrilla de San Martin et al., 2020) and in the SST-expressing, Oriens Lacunosum-Moleculare (OL-M) INs to PN synapse (Schulz et al., 2018). Furthermore, in the rat somatosensory cortex, PV-IN-mediated PN perisomatic inhibition is sensitive to zolpidem (100 nM), a positive allosteric modulator, which at this concentration, is known to specifically bind the benzodiazepine site of α1-containing and, less efficiently, α2- and α3-containing GABAA receptors (Korpi et al., 2002; Möhler, 2002; Bacci et al., 2003). In order to validate these results in the mouse somatosensory cortex we tested the effects of α5IA, a negative allosteric modulator (NAM) specific for α5-GABA_A_Rs (Dawson et al., 2006), and zolpidem on both MC-mediated PN dendritic inhibition and PV-IN-mediated PN perisomatic inhibition (Fig 4 A). PV-PN uIPSC weighted decay time constant (τd,w) was significantly increased by zolpidem (control: 9.0 ± 1.3 ms; zolpidem: 11.2 ± 0.7 ms, n=6 pairs, p=0.014, Paired t-test; Fig 4 B). In contrast, PV-PN uIPSCs amplitude was unaffected by α5IA (control: 63 ± 22 pA; α5IA: 65 ± 20 pA, n=6 pairs, p=0.7294, Paired t-test; Fig. 4 B). The amplitude of uIPSCs elicited from MCs were highly sensitive to α5IA (control: 177 ± 44 pA; α5IA: 104 ± 23 pA, n=11 pairs; p=0.003, Wilcoxon Signed-Ranks test; Fig 4 C), and zolpidem did not affect the weighted decay time constant of the MC-PN uIPSCs (control: 8.2 ± 1.2 ms; zolpidem: 9.1 ± 0.9, n=6 pairs, p=0.173, paired T test, Fig 4 C). Importantly, α5IA is a partial NAM displaying ~40% efficacy, thus not providing a complete blockade of α5-GABA_A_Rs (Dawson et al., 2006). Importantly, after incubation with α5IA, the remaining MC-PN uIPSC amplitude was near 60% (65.7% ± 5.4%; Fig. 4 C). This suggests that unitary synaptic responses from MCs to PNs are fully mediated by α5-GABA_A_Rs.

**Figure 4:**
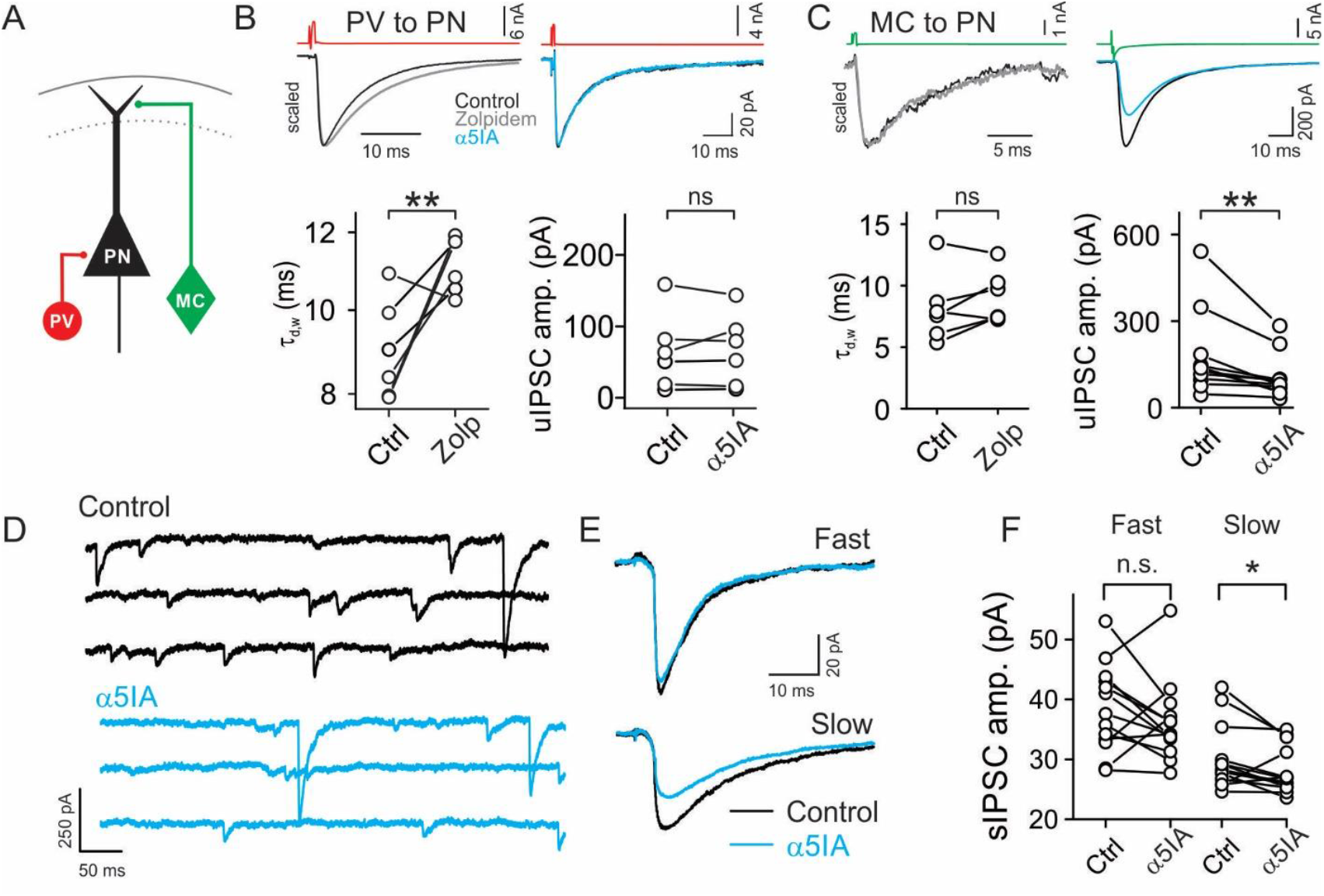
α5-GABA_A_Rs mediate synaptic inhibition selectively from MCs. **A:** Schematic of paired recordings between a MC or PV-IN and a PN. **B:** Top left, Representative average uIPSC elicited by a PV-INs onto a PN in the absence (black) and presence (grey) of zolpidem. Traces are scaled to highlight zolpidem effect on uIPSC decay time. Bottom left, population data of zolpidem effect on the weighted decay time constant (τ_d,w_, left) and α5IA effect on uIPSC amplitude in PV-PN pairs. Top right: representative average uIPSC traces elicited a PV cell onto a PN in the absence (black) and presence (blue) of α5IA. **C:** Same as in B for MC-PN pairs. **D:** Representative voltage-clamp traces of sIPSCs recorded from a PN before (control, black) and after 15 minutes incubation with 100 nM α5IA (blue). **E:** Representative averaged traces of fast (top) and slow (bottom) events recorded from MCs. Only amplitudes of slow events are affected by α5IA (blue trace, bottom panel). **F:** Population plot of individual cells, fast and slow sIPSCs median amplitudes measured in control and after incubation with α5IA. *p<0.05, **p<0.01.

α5-GABA_A_Rs have been hypothesized to be extrasynaptic, mainly mediating tonic inhibition (Caraiscos et al., 2004). However, there is growing evidence that α5-GABAAR are also involved in dendritic inhibition at specific synapses made by MCs in the cortex and by OLM interneurons in the hippocampus (Ali and Thomson, 2008; Schulz et al., 2018; Zorrilla de San Martin et al., 2020). It is possible that sensitivity of uIPSCs to α5IA could be partially or fully due to activation of extrasynaptic GABA_A_Rs due to GABA spillover, induced by AP-evoked synaptic transmission. To further study the role of synaptic α5-GABA_A_Rs, we measured spontaneous inhibitory postsynaptic currents (sIPSCs) recorded from PNs (Fig 4 D-F). Because quantal, AP-independent synaptic events make up a large fraction of sIPSCs, these are less likely shaped by activation of extrasynaptic receptors. To separate putative dendritic and perisomatic events, we sorted sIPSCs into two groups based on their rise-times (Fig 4 D-F). We considered the events with rise times larger than 1.8 ms as ‘slow’, whereas those with rise-times smaller than 1.8 ms were defined as ‘fast’, based on the average rise-time obtained at connected MC-PN pairs (Figs. 1I; 2C-E). The amplitudes of slow sIPSCs were significantly reduced after 10 minutes incubation with 100 nM α5IA (control: 31 ± 2 pA, α5IA: 28 ± 1 pA, n=11 cells, p=0.03, Wilcoxon Signed-Ranks test; Fig. 4 E). Conversely, the same concentration of α5IA did not affect fast sIPSCs amplitude (control: 38 ± 2 pA, α5IA: 36 ± 2 pA; n=11 cells, p=0.3636, Wilcoxon Signed-Ranks test; Fig 4 E). This result indicates that fast, perisomatic events are generated by other interneurons types, not using α5-GABA_A_Rs.

Altogether, these results indicate that synapses formed by dendrite-targeting MCs onto PNs, specifically express α5-GABA_A_Rs whereas PV-PN synapses express α1-GABA_A_Rs.

### MCs inhibit PNs, but not other interneurons, through α5-GABA_A_Rs

In the previous sections, we showed that MCs make synaptic contacts exhibiting target-specific biophysical and physiological properties. We also showed that, among the inhibitory inputs received by PNs, those originating from MCs are distinguished by their sensitivity to α5IA. We therefore tested whether postsynaptic expression of α5-GABA_A_Rs is a trait shared by all synaptic contacts made by MCs or it is specific for synaptic contacts that MCs form on PN dendrites. To address this question, we measured unitary GABAergic synaptic transmission between MCs and other INs and tested their sensitivity to α5IA. We found that uIPSC amplitudes elicited by MCs and recorded in PV INs (control: 31 ± 4 pA, α5IA: 35 ± 6 pA, n=11 pairs, p=0.3757, paired t test), L1 INs (control: 83 ± 32 pA, α5IA: 102 ± 29 pA, n=5 pairs, p=0.3757, paired t test) and VIP INs (control: 178 ± 100 pA, α5IA: 118 ± 52 pA, n=4 pairs, p=0.7432, Wilcoxon signed ranks test) were not sensitive to α5IA (Fig. 5 B-C).

**Figure 5:**
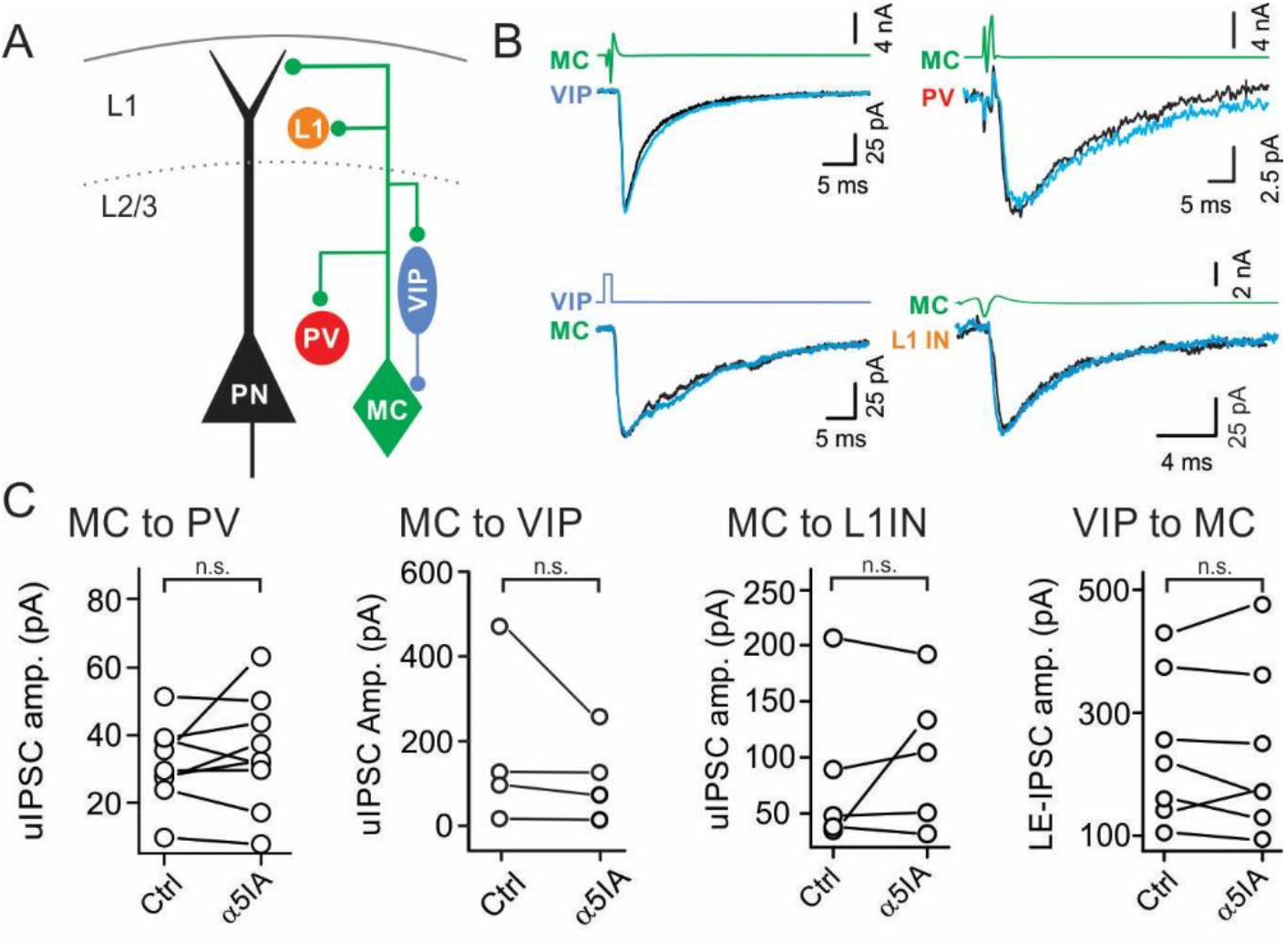
Inhibitory synaptic transmission involving MCs and other interneurons does not rely on α5-GABA_A_Rs. **A:** Schematic representation of the tested inhibitory circuits involving MCs. **B:** Representative averaged voltage-clamp traces of uIPSCs from MCs onto different element of the circuit and from VIP to MC before (black) and after (blue) application of α5IA. **C:** Population data of uIPSC amplitude before (Ctrl) and 15 minutes after α5IA application.

In the hippocampus, SST-positive, OL-M INs receive α5-mediated inhibition from VIP INs (Magnin et al., 2019). Since VIP-MC synapses represent an important disinhibitory circuit in the cortex as well, we asked whether α5-GABA_A_Rs mediate inhibitory inputs from VIP INs also in the mouse somatosensory cortex. To address this question and to activate VIP INs specifically, while recording from GFP-expressing MCs, we crossed VIP-cre mice with X98 mice. Since dual whole-cell recordings showed a very low yield (3 connected out of 45 recorded pairs), we expressed the light-sensitive opsin ChR2 via injection of flexed-ChR2 AAV particles in the barrel cortex of VIPCre::X98 1-3-days-old pups. We recorded light-evoked IPSCs in MCs, and found that inhibitory responses originating at VIP cells were not sensitive to α5IA (control: 170 ± 46 pA, α5IA: 166 ± 52 pA, n=7, p=0.7432, Wilcoxon signed ranks test; Fig 5 B). Furthermore, the amplitude of sIPSCs recorded from MCs were not affected by incubation with 100 nM α5IA (control: 32 ± 1 pA, α5IA: 29 ± 3 pA, n=26, p=0.0659, Wilcoxon signed ranks test; Fig 5 E).

Overall, these results indicate that GABAergic inhibition to and from MCs uses α5-GABA_A_Rs exclusively at synapses formed with PN distal dendrites and not for other MC-targets within cortical circuits. Thus, the characteristic slow kinetics of uIPSCs, the distal dendritic targeting and the synaptic expression of α5-GABA_A_Rs represent unique molecular and cellular signatures of MC-PN synapses.

### Tonic inhibition is mediated by α5-GABA_A_Rs in PN, but not MC nor PV-IN

α5-GABA_A_Rs have been largely associated to tonic inhibition due to extrasynaptic immunoreactivity in cell culture (Loebrich et al., 2006; Serwanski et al., 2006), hippocampus and cortex (Serwanski et al., 2006) and amygdala (Botta et al., 2015) and the lack of tonic inhibitory current in hippocampal PNs of α5 knock-out mice (Caraiscos et al., 2004).

Thus, we pre-incubated slices with ACSF or ACSF + 100 nM α5IA during at least 10 minutes and then quantified the difference in holding current amplitude (ΔI_hold_) before and after bath application of 1 μM gabazine (Fig. 6). Pre-incubation with 100 nM α5IA significantly reduced GABAergic ΔIhold in PNs as compared to slices preincubated in ACSF only (ACSF: 51 ± 1 pA, n=16; α5IA: 29 ± 7 pA, n=23, p=0.0244, unpaired t test; Fig. 6 A,B). A similar percentage of reduction was obtained incubating with 500 nM α5IA (data not shown), ruling out that α5-GABA_A_Rs required higher drug concentrations. Conversely, incubation with α5IA failed to produce any significant change in tonic current recorded from MCs (ACSF: 16 ± 5 pA, n=9; α5IA: 23 ± 6 pA, n=14, p=0.3330, unpaired t test; Fig. 6 C,D) nor PV INs (ACSF: 33 ± 10 pA, n=9; α5IA: 24 ± 5 pA, n=14, p=0.6591, unpaired t test; Fig. 6 E,F), showing that tonic inhibition is mediated by α5-GABA_A_Rs exclusively in PNs.

**Figure 6:**
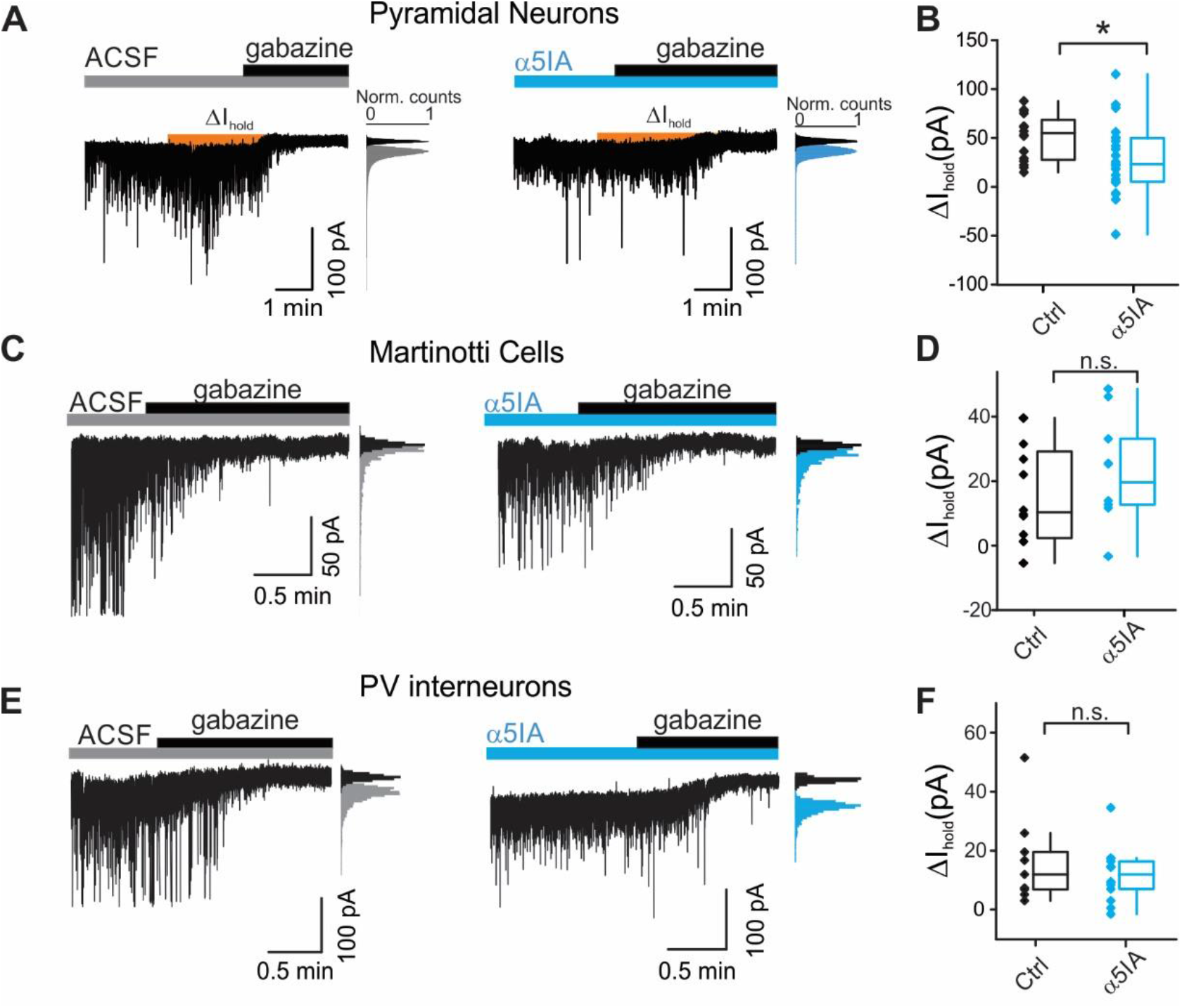
α5GABA_A_Rs only contribute to tonic inhibition in L2/3 PN of mouse somatosensory cortex. **A:** Whole-cell voltage-clamp recordings from L2/3 PNs preincubated with either vehicle (aCSF, left) or α5IA (right). DNQX (10 μM) and GABA (5 μM) were continuously present in both conditions. Orange areas (ΔIhold) represent tonic inhibition measured after gabazine onset (dotted red line). Insets: All-points histograms of the current trace obtained in the absence (grey and blue histograms) and presence of gabazine (black histograms). Gaussian fits were used to determine the noise half-width. **C-E:** same as in A for MCs and PV-INs, correspondingly. **B-D-F:** Population graphs of holding-current shifts after gabazine application (ΔI_hold_). * p<0.05.

Altogether, these results indicate that α5-GABA_A_Rs are selectively expressed by PNs and they mediate both tonic and dendritic, phasic synaptic inhibition.

## Discussion

In this study, we explored physiological aspects of different inhibitory synaptic connections made by MCs in the L2/3 of the mouse somatosensory cortex. We show that inhibitory synapses made by MCs display biophysical, morphological and pharmacological properties that are specific in distinct postsynaptic partners. We found that the most extensively contacted cells are PNs. However, also PV, VIP and L1 inhibitory INs also receive significant inhibition from MCs. Notably, we showed that the MC-PN synapse is distinguished by two unique features: slow kinetics and expression of the α5-GABAAR. Finally, we showed that α5-GABA_A_Rs contribute to tonic inhibition of PNs but not of other INs, confirming the specific involvement of this receptor in PN dendritic inhibition.

SST-cre mouse lines are widely used to study the functional role of SST-expressing INs by manipulating and recording their activity, using cre-driven expression of light-sensitive opsins or genetically encoded Ca^2+^ sensors (Taniguchi et al., 2011). Despite its extensive use, the SST-cre mouse line affects all interneurons expressing SST. Yet, SST-positive neurons encompass an heterogeneous group of inhibitory neurons that display different morphology and spiking patterns as well as diverse connectivity (Halabisky et al., 2006; Ma et al., 2006; McGarry et al., 2010; Naka et al., 2019). Therefore, it is important to focus the study of SST-positive neurons to defined homogeneous subpopulations, in order to prevent unwanted over-simplified conclusions. MCs represent a specific subpopulation of GABAergic interneurons, accounting for only 20% of all SST-expressing cells (Yavorska and Wehr, 2016). Here we used the X98 mouse line to study MCs specifically (Ma et al., 2006). We provide evidence that GFP-expressing neurons in the somatosensory cortex from the X98 mouse line exhibit the typical anatomical and electrophysiological properties of MCs (Wang et al., 2004; Kapfer et al., 2007; Silberberg and Markram, 2007; Tremblay et al., 2016). In addition, glutamatergic recruitment of GFP-positive cells is strongly facilitating, as opposed to PV cells, another hallmark of MCs (Reyes et al., 1998). Therefore, we conclude that this mouse line represents an excellent tool to study inhibitory circuits involving MCs.

Even though MCs extensively inhibit PNs via α5-GABA_A_Rs, they also contact other elements of the cortical microcircuits, and, in addition, they are targeted by VIP-expressing interneurons (Pfeffer et al., 2013; Kepecs and Fishell, 2014; Tremblay et al., 2016; Walker et al., 2016). We found that MCs contact PV-VIP-and L1-INs at a reduced connectivity rate, as compared to MC-PN connections. Moreover, GABAergic synapses from MCs onto other interneurons and those inhibiting MCs from VIP-INs do not use α5-GABA_A_Rs.

MCs were hypothesized to provide a non-specific ‘blanket’ of inhibition to PNs (Fino and Yuste, 2011; Fino et al., 2013; Karnani et al., 2016). Accordingly, we found a relatively high connectivity rates between MCs and L2/3 PNs, consistent with the prominent MC axonal plexus innervating L1. However, our results indicate that despite extensively innervating L1, MC axons possess a very strong tropism for PN dendrites. Yet, despite at lower rate, MCs do contact also L1-INs, which exert slow feed-forward inhibition on PN dendrites during the encoding of context-rich, top-down information from higher order thalamus and cortices (Letzkus et al., 2011; Abs et al., 2018).

Importantly, dendritic inhibition seems to be a specific feature of MC-PN connections, as uIPSC rise times measured on other MC targets (interneurons) had fast (<1 ms) kinetics similar to the known PV-PN perisomatic responses. In agreement with this view, we failed to find evidence of direct contact between MC axons on the perisomatic region of PNs. Conversely, we found MC putative boutons juxtaposed to the perisomatic region of PV, VIP and L1 INs. This is consistent with the fast, non-filtered, IPSCs observed in somatic whole-cell recordings and in line with a previous report showing that inhibitory contacts onto PV INs are preferentially located in the proximal dendrites and soma while excitatory inputs are located in distal dendrites (Kameda et al., 2012).

Use-dependent short-term facilitation or depression of synaptic responses has been traditionally linked to presynaptic loose- or tight-coupled synapses, identifying diverse cell types with specific biophysical presynaptic properties, such as low or high release probability, respectively (Jackman and Regehr, 2017). Importantly, frequency-dependent bidirectional short-term plasticity is a powerful synaptic tool to provide distinct cell types with a specific strategy to transfer information about presynaptic spike trains. We found that GABAergic synapses from MCs exhibit a stark target-cell-specific facilitation and depression. Target-cell-specific short-term plasticity and release probability, originating from the same cell type, was described at glutamatergic synapses from PNs recruiting different IN subtypes in the neocortex and hippocampus (Reyes et al., 1998). Our finding indicates that single-axon, target-specific bidirectional short-term plasticity occurs also at GABAergic synapses. Intriguingly, synapses made in L1 (with either PN distal dendrites or sparse INs) are depressing, whereas, inhibitory connection that the same cells make onto their targets in L2/3 (PV and VIP cells) are either uniform or strongly facilitating. It will be interesting to determine the molecular and synaptic mechanisms by which the identity of the postsynaptic neuron determines the efficacy of GABAergic synapses originating from the same MC. Target cell type-dependent variability in presynaptic properties increases the computational power of neuronal networks. It will be therefore fundamental to understand the functional role of such a target-specific regulation of inhibitory synaptic efficacy.

MCs exhibit differential inhibitory strategies depending on the postsynaptic cell type: they modulate input onto PNs and they likely control, at least in part, the output activity of other interneurons. Therefore, inhibitory circuits formed by MCs seem to exhibit a more complex architecture and function than previously hypothesized as provider of a mere blanket of inhibition (Fino and Yuste, 2011).

In addition to the strong preference for distal apical dendrites, MCs display another PN-specific synaptic feature, as they use α5-GABA_A_Rs for synaptic dendritic inhibition. Indeed, GABAergic synapses from MCs to other interneurons are perisomatic and do not use α5-GABA_A_Rs. In fact, lack of α5IA effects on tonic inhibition on PV and MCs suggest that these major IN subtypes do not express this GABAAR α subunit. Interestingly, It has been recently reported that hippocampal Oriens-Lacunosum Moleculare INs also express, α5-GABA_A_Rs at synapses originating at VIP INs (Magnin et al., 2019). Yet, we did not find evidence of α5IA effect on VIP-IN-evoked IPSCs in MCs of the barrel cortex, suggesting that cortical MCs differ from their hippocampal counterparts. The α5 subunit is much more strongly expressed in the hippocampus than in the neocortex (Lingford-Hughes et al., 2002). Therefore, it will be interesting to reveal whether α5 has different circuit-specificity and/or plays a different role in these two areas. Likewise, it remains to be tested whether α5-GABA_A_Rs are also expressed by other subtypes of inhibitory neurons. Our results on L1-INs suggest that MCs do not use α5-GABA_A_Rs at these synapses. However, L1 is populated by a heterogeneous IN population (Schuman et al., 2019) and, since we did not use specific mouse lines to target distinct cell types, our data may have been collected from a relatively heterogeneous interneuron group.

In addition to dendritic filtering, MC-PN synaptic responses might be slow due to the specific properties of the α5-subunit itself, which is exclusively expressed at this synapse. The slow kinetics and the rectification of α5-GABA_A_Rs match the biophysical properties of NMDARs, which govern Ca^2+^ signaling and dendritic computation in PNs (Branco and Häusser, 2010; Tran-Van-Minh et al., 2015; Schulz et al., 2018). Dendritic patch would be necessary to test this hypothesis, although the high series resistance typical of dendritic patch recordings might prevent an accurate analysis of fast currents.

α5-GABA_A_Rs have been proposed to mediate tonic inhibition due to their sensitivity to nanomolar concentrations of GABA, their non-desensitizing properties and the lack of evidence supporting its implication in synaptic transmission (Caraiscos et al., 2004). However, knock down of radixin, the extrasynaptic scaffolding protein associated to α5-GABA_A_Rs did not produce any effect on GABA evoked current, suggesting that extrasynaptic α5-GABA_A_Rs might not be functional (Loebrich et al., 2006). Furthermore, the participation of α5-GABA_A_Rs in phasic synaptic inhibition has been recently demonstrated in different brain structures, namely the rat somatosensory cortex (Ali and Thomson, 2008), mouse hippocampus (Schulz et al., 2018) and mouse prefrontal cortex (Zorrilla de San Martin et al., 2020). Even for action potential-dependent unitary responses between MCs and PNs, it is possible that GABA could spill over to peri- or extrasynaptic GABA_A_Rs containing α5. If this were the case, we would not have detected significant effects on quantal events, which reflect mostly purely synaptic activation of GABA_A_Rs. Importantly, we recorded sIPSCs from the soma of L2/3 PNs and found that only slow sIPSCs were sensitive to α5IA, whereas fast perisomatic inhibitory events were unaffected. Our results on sIPSCs corroborate the synaptic localization of α5-GABA_A_Rs. Indeed, at our extracellular K^+^ concentrations, sIPSCs are dominated by AP independent miniature events. The blockade of MC-PN uIPSCs, slow sIPSCs and tonic inhibition was not total but it was in all cases maximal, taking into account the actual efficacy (~40%) of α5IA (Sternfeld et al., 2004; Atack, 2010).

Therefore, the most parsimonious interpretation of our pharmacological experiments is that α5-GABA_A_Rs are prominently expressed at synaptic sites of dendritic MC-PN connections and are responsible for dendritic inhibition from this specific GABAergic neuron type. In fact, the α5-mediated tonic currents could be the direct activation by ambient GABA of high affinity synaptic, and not necessarily extrasynaptic receptors.

The specific expression of the α5 GABAAR subunit in PNs is particularly interesting in light of its involvement in cognitive processes. Mice lacking the *Gabra5* gene, encoding for the α5 subunit of the GABAAR, show enhanced performance in cognitive tasks (Collinson et al., 2002). This evidence, in addition to the high α5-GABA_A_Rs expression in the mouse prefrontal cortex and hippocampus (Fritschy and Mohler, 1995) led to propose novel potential pro-cognitive pharmacological strategies. This strategy is being actively explored to treat intellectual disability in Down syndrome (Braudeau et al., 2011; Martínez-Cué et al., 2013; Duchon et al., 2019; Zorrilla de San Martin et al., 2020) and in other brain diseases characterized by memory impairments (Zurek et al., 2014) and depressive states (Zanos et al., 2017). Specific negative modulation of these receptors would facilitate cognition avoiding anxiogenic and pro-convulsive effects of wide spectrum GABA_A_Rs antagonists due to the restricted expression of the α5 subunit to this specific inhibitory circuit formed by MCs.

## Acknowledgments

We thank the ICM technical facilities PHENO-ICMICE, iGENSEQ and ICM.Quant. This work was supported by “Investissements d’avenir” ANR-10-IAIHU-06, BBT-MOCONET1 ; BBT-MOCONET2; Agence Nationale de la Recherche (ANR-13-BSV4-0015-01 ; ANR-16-CE16-0007-02 ; ANR-17-CE16-0026-01 ; ANR-18-CE16-0011-01 ; ANR-20-CE16-0011-01; ANR-12-EMMA-0010;), Fondation Recherche Médicale (Equipe FRM DEQ20150331684 and EQU201903007860), NARSAD independent Investigator Grant and Fondation Lejeune (#1790). CD was supported by the École de Neurosciences de Paris et Île de France (ENP) and by the Labex Bio-Psy. PHENO-ICMICE was supported by two “Investissements d’avenir” (ANR-10-IAIHU-06 and ANR-11-INBS-0011-NeurATRIS) and the “Fondation pour la Recherche Médicale”.

